# Host genotype and stable differences in algal symbiont communities explain patterns of thermal stress response of *Montipora capitata* following thermal pre-exposure and across multiple bleaching events

**DOI:** 10.1101/2020.07.24.219766

**Authors:** Jenna Dilworth, Carlo Caruso, Valerie A. Kahkejian, Andrew C. Baker, Crawford Drury

## Abstract

As sea surface temperatures increase worldwide due to climate change, coral bleaching events are becoming more frequent and severe, resulting in reef degradation. Leveraging the inherent ability of reef-building corals to acclimatize to thermal stress via pre-exposure to protective temperature treatments may become an important tool in improving the resilience of coral reefs to rapid environmental change. We investigated whether historical bleaching phenotype, coral host genotype, and exposure to protective temperature treatments would affect the response of the Hawaiian coral *Montipora capitata* to natural thermal stress. Fragments were collected from colonies that demonstrated different bleaching responses during the 2014-2015 event in Kāne’ohe Bay (O’ahu, Hawai’i) and exposed to four different artificial temperature pre-treatments (and a control at ambient temperature). After recovery, fragments experienced a natural thermal stress event either in laboratory conditions or their native reef environment. Response to thermal stress was quantified by measuring changes in the algal symbionts’ photochemical efficiency, community composition, and relative density. Historical bleaching phenotype was reflected in stable differences in symbiont community composition, with historically bleached corals containing only *Cladocopium* symbionts and historically non-bleached corals having mixed symbiont communities dominated by *Durusdinium*. Mixed-community corals lost more *Cladocopium* than *Cladocopium*-only corals during the natural thermal stress event, and preferentially recovered with *Durusdinium*. Laboratory pre-treatments exposed corals to more thermal stress than anticipated, causing photochemical damage that varied significantly by genotype. While none of the treatments had a protective effect, temperature variation during treatments had a significant detrimental effect on photochemical efficiency during the thermal stress event. These results show that acclimatization potential is affected by fine-scale differences in temperature regime, host genotype, and relatively stable differences in symbiont community composition that underpin historical bleaching phenotypes in *M. capitata*.

## Introduction

Tropical coral reefs are one of the most biodiverse and ecologically valuable ecosystems on the planet (Connell 1978). Worldwide, reefs are in decline due to a combination of stressors (Hughes et al. 2003; Pandolfi et al. 2003) including climate change, which is leading to rising sea surface temperatures and ocean acidification (Hoegh-Guldberg et al. 2007). Bleaching induced by these increased temperatures is a severe threat to the persistence of coral reef ecosystems: the coral’s obligate endosymbiosis with dinoflagellate algae in the family Symbiodiniaceae (LaJeunesse et al. 2018) can break down under thermal stress due to the dinoflagellate’s increased production of oxidative agents (Glynn 1993; Lesser 2006; Weis 2008). Tropical reef-building corals live within 1-2°C of their upper thermal limits (Coles et al. 1976), so rising sea surface temperatures and more frequent temperature anomalies have lead to increasingly frequent and more severe coral bleaching events worldwide in recent decades (Hoegh-Guldberg et al. 2007). Annual bleaching is expected to impact most of the world’s coral reefs by mid-century if global carbon emissions continue along the IPCC’s RCP8.5 business-as-usual trajectory (van Hooidonk et al. 2013; van Hooidonk et al. 2014).

However, the resistance and response of reef-building corals to bleaching is variable within and between species (Glynn 1996; Cunning et al. 2016; Schoepf et al. 2019), indicating an ability to acclimatize and adapt to increasing thermal stress. Adaptation results in gradual adjustment to changing environmental conditions at the population level (van Oppen et al. 2015), but corals will likely only be able to successfully adapt to a changing climate if current greenhouse gas emissions are significantly reduced (Császár et al. 2010). The adaptation rate of scleractinian corals is likely too slow to keep pace with the rapid environmental changes expected under future climate change (Bay et al. 2017; Matz et al. 2018), and the adaptive capacity of many coral reefs based on standing diversity is expected to be overwhelmed within a century (Matz et al. 2020). Acclimatization occurs through non-genetic changes at the individual rather than population level and can take place more rapidly, within a single generation (van Oppen et al. 2015). Some mechanisms of acclimatization, such as non-coding epigenetic changes to DNA through methylation or chromatin modification (van Oppen et al. 2015), may be passed on to offspring and thus change the response of future generations to thermal stress (Putnam and Gates 2015; Torda et al. 2017). The intra- and intergenerational acclimatization of corals may interact to bridge the gap between current populations in a rapidly warming environment and the ability of corals to adapt to these changing conditions (Chevin et al. 2010; Palumbi et al. 2014; Drury 2019).

Previous work shows that a history of exposure to thermal stress can affect the response of corals to bleaching events (Maynard et al. 2008) even without genetic changes (Brown et al. 2015), indicating the involvement of acclimatization mechanisms. In several lab-based experiments, pre-exposing corals to increased temperatures before thermal stress improved subsequent thermal tolerance (Middlebrook et al. 2008; Bellantuono et al. 2012a, 2012b; Bay and Palumbi 2015; Ainsworth et al. 2016), indicating that acclimatization to thermal stress can be induced. In these studies, increased thermal tolerance was underpinned by changes in host gene expression (Bellantuono et al. 2012b; Bay and Palumbi 2015; Ainsworth et al. 2016) and the physiological response of algal symbionts (Middlebrook et al. 2008), rather than shifts in the composition of the algal symbiont community to more thermally tolerant symbionts, known as symbiont “shuffling” (Baker 2003; Silverstein et al. 2014; Cunning et al. 2015).

Given the key role acclimatization to thermal stress will likely play in the survival of coral reefs under rapidly changing environmental conditions, it is important to better understand the potential and limits of utilizing artificially induced acclimatization to increase coral resilience to thermal stress. In this study, we explore the utility of artificial temperature profiles in a laboratory setting to induce acclimatization and the response of pre-treated corals to a natural thermal stress event using *Montipora capitata* in Kāne’ohe Bay (O’ahu, Hawai’i). *M. capitata* is one of the dominant reef builders in Kāne’ohe Bay (Bahr et al. 2015a) and harbors symbionts of the genera *Cladocopium* (bleaching susceptible) and/or *Durusdinium* (bleaching resistant) (Rowan 2004; Cunning et al. 2016; Innis et al. 2018). During the 2015 bleaching event, immediately adjacent pairs of *M. capitata* colonies where one colony bleached and the other did not were tagged based on their observed bleaching phenotype (Cunning et al. 2016; Matsuda et al. 2020) A subset of these tagged colonies represent distinct genotypes (Drury unpublished data). Using this unique framework of known historical bleaching phenotype and host genotype, we 1) examine the effectiveness of different temperature profiles in inducing an acclimatization response in *M. capitata*, 2) determine whether the effects of these profiles can persist throughout the natural warming season in Kāne’ohe Bay, and 3) examine the contribution of temperature treatment, host genotype, and historical phenotype to the thermal stress response. This work provides insight into the potential for using artificially induced acclimatization to increase thermal tolerance in reef-building corals, as well as elucidating how genotypic and phenotypic differences may influence the response of temperature pre-treated corals to thermal stress.

## Materials and Methods

### Coral fragment collection and acclimation

*M. capitata* fragments were collected on 11^th^ June 2019 from patch reef 13 in Kāne’ohe Bay, O’ahu, Hawai’i (Figure 1a). We collected 50 fragments each from 10 tagged colonies of unique genotype and known bleaching phenotype in the 2015 event (five non-bleached, five bleached), in addition to five larger sacrificial fragments from each colony intended for tissue sampling, resulting in 500 small fragments and 50 large fragments. The tagged colony framework minimized environmental differences between colonies, allowing us to focus on genotypic and phenotypic differences. All corals were visibly healthy at the start of the experiment, and are thought to be physiologically and reproductively recovered from the 2015 bleaching event (Cunning et al. 2016; Wall et al. 2019). Coral fragments were mounted on aragonite plugs and allowed to acclimate to lab conditions for four weeks. The fragments were allowed to recover in outdoor water tables at ambient temperature, after which they were transferred to indoor tanks for the final week of acclimation on a 12:12hr light:dark cycle with a maximum midday irradiance of ∼383 μmol/sec at ambient temperature before beginning the experiment.

**Fig. 1.**
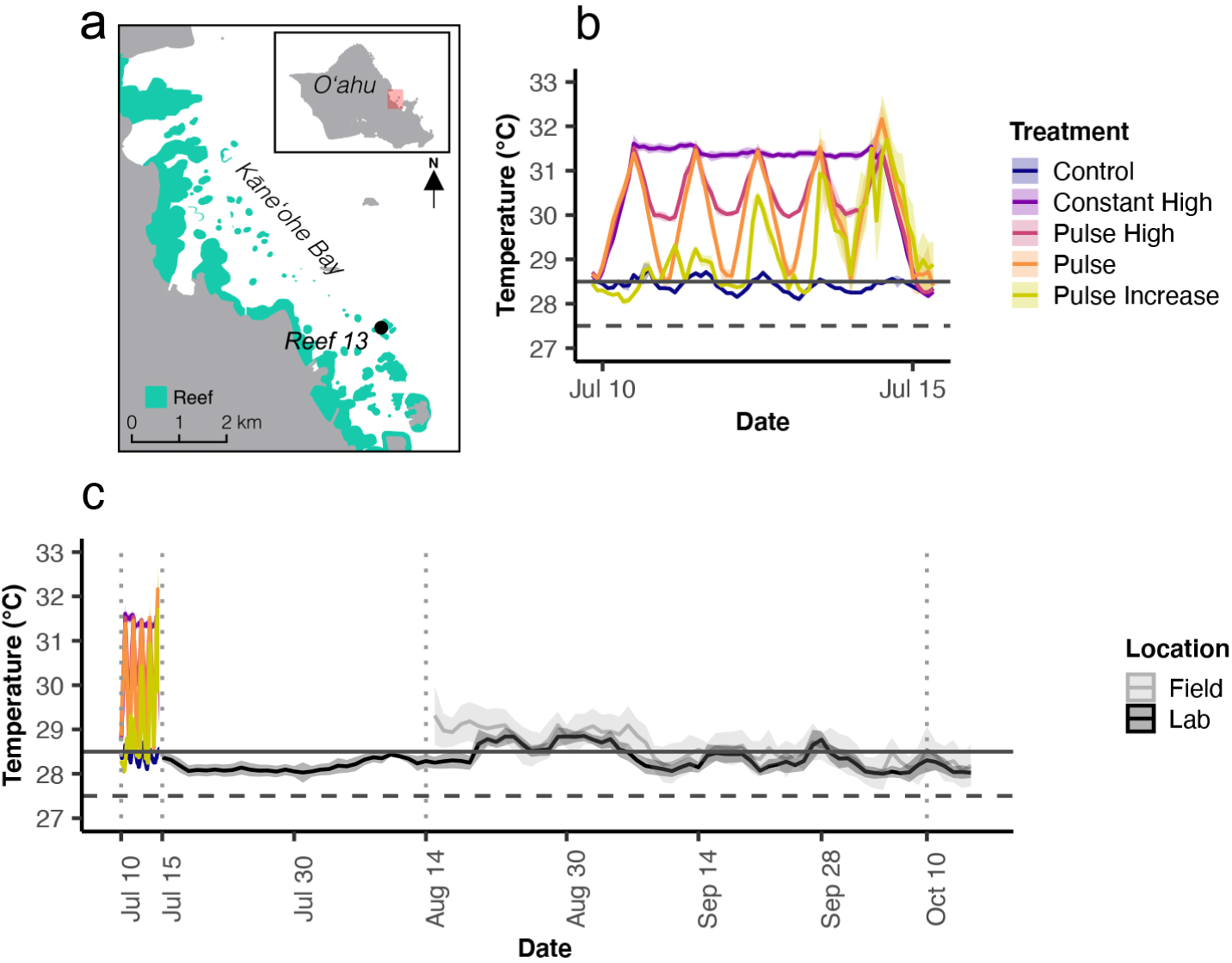
**(a)** Map showing the location of patch reef 13 in Kāne’ohe Bay, O’ahu, Hawai’i. Areas shaded in green indicate fringe/patch reefs. **(b)** Mean hourly temperatures in the four temperature treatments and control. Colors indicate the five treatments: control (blue), constant high (purple), pulse high (pink), pulse (orange), and pulse increase (yellow). Shaded areas represent the standard deviation. The dashed horizontal line indicates the maximum monthly mean for Kāne’ohe Bay (27.5°C), while the solid horizontal line indicates the bleaching threshold (28.5°C). **(c)** Temperatures throughout the experiment. July 9^th^ – July 15^th^: mean hourly temperatures in the four acclimatization treatments and control. Colors the same as panel (a). July 16^th^ – October 20^th^: mean daily temperatures in the lab (black) and field (grey). Shaded areas represent the standard deviation. Dotted vertical lines indicate sampling timepoints for PAM fluorometry and tissue sampling. Temperature treatments took place in the time period within the first two vertical lines. The natural thermal stress event occurred between the second two vertical lines. The dashed horizontal line indicates the maximum monthly mean for Kāne’ohe Bay (27.5°C), while the solid horizontal line indicates the bleaching threshold (28.5°C).

### Measures of acclimatization and thermal stress response

Pulse amplitude modulation (PAM) fluorometry was used to measure the dark-adapted quantum yield (F_v_/F_m_) of photosystem II, which has been used extensively as a measure of damage to photosystem II in the algal symbiont and thus as an indicator of thermal stress and bleaching (Warner 1999; Jones et al. 2000). A diving-PAM II (Walz) was used to monitor the 500 small coral fragments throughout the experiment. Additionally, tissue samples were taken from the sacrificial large fragments at key timepoints throughout the study: pre-treatment, post-treatment, after treatment recovery, and at the peak of the natural thermal stress event (Figure 1c). At the natural thermal stress timepoint, a subset of 200 small fragments were also sampled. Samples were taken from the midpoint of the branch using a 5mm diameter dermal curette, taking care to avoid previously sampled tissue. Tissue samples were kept on ice until all fragments had been sampled, then frozen at -80°C for DNA extraction and molecular analysis of symbiont to host cell ratio and symbiont community composition using quantitative PCR (qPCR).

### Temperature profiles

Four different artificial temperature profiles were developed (Figure 1b). Based on preliminary data (see the Supplementary Materials), 31.5°C was chosen as the maximum temperature to ensure coral fragments in all treatments, including constant high temperature conditions, would be exposed to sublethal thermal stress. Pulsed temperature profiles were successfully used by Bay and Palumbi (2015) to elicit an acclimatization response in the coral host, so we compared various pulsed profiles to constant temperatures using four treatments: pulse, pulse high, pulse increase, and constant, in addition to a control at ambient temperature (Figure 1b). The five treatments were randomly assigned to duplicate tanks and fragments were randomized so that five fragments from each of the 10 genotypes were assigned to each tank. Larger fragments were randomized so that one fragment from each genotype was assigned to one of the five treatments. Temperatures in the tanks were recorded throughout the experiment using HOBO Pendant temperature loggers (Onset Computer Corporation). During the temperature treatments, fragments were monitored daily with PAM fluorometry, and tissue samples were taken from the large fragments before and after the temperature treatments (Figure 1c). After five days, fragments began to start showing a decline in F_v_/F_m_, so temperatures were ramped back down to ambient temperatures, as the goal of the treatment period was to expose coral fragments to protective levels of stress only. Following the treatments, fragments were allowed to recover in their tanks at ambient temperature for four weeks. Tissue samples and PAM fluorometry measurements were taken at the end of the recovery period (Figure 1c).

### Natural thermal stress event

Following the four-week recovery period, fragments were split into two groups: 200 small fragments and the 50 larger fragments were kept in the lab while the remaining 300 fragments were returned to the field on 15^th^ August 2019 in anticipation of a bleaching event in Kāne’ohe Bay (Figure 1c). Coral fragments were randomized such that three fragments per genotype in each tank were returned to the reef, and two fragments per genotype per tank remained in the lab. Corals deployed into the field were attached to a single table at reef 13 in Kāne’ohe Bay with triplicate temperature loggers. Field corals were returned to the lab on 10^th^ October 2019 for PAM fluorometry with the fragments that remained in the lab. Tissue samples were taken from the large fragments and from a subset of 200 (four fragments per colony per treatment) of the corals that were deployed to the reef. While water temperatures in the lab were slightly lower than those in the field, temperatures in both locations tracked similar trajectories, and corals in both conditions were exposed to temperatures above the bleaching threshold of 28.5°C (Figure 1c; NOAA/NDBC; Wyatt et al. 2019). The 10^th^ October 2019 sampling timepoint for PAM fluorometry and tissue samples was considered the peak of thermal stress, as the water temperature maximum had already occurred (Figure 1c).

### Molecular analyses

In December 2019 and January 2020, DNA was extracted from tissue samples taken at the four key timepoints (Figure 1c) with a CTAB-chloroform protocol (dx.doi.org/10.17504/protocols.io.dyq7vv). Concentration of *Cladocopium, Durusdinium*, and coral host cells in these samples was quantified with qPCR using actin and Pax-C assays (Cunning and Baker 2013; Cunning et al. 2015; Cunning et al. 2016) on a StepOnePlus system (Applied Biosystems) with two technical replicates and a fluorescence threshold of 0.01 run for 40 cycles. Baseline values were selected on amplification cycle threshold (C_T_) values (around 5- 23). C_T_ values were corrected for fluorescence and copy number using the equations of Cunning, Ritson-Williams, and Gates (2016). Samples were re-extracted and/or re-analyzed if C_T_ replicate standard deviation was greater than one between replicates, or if only a single replicate amplified.

### Statistical analyses

Following quality control (see the Supplementary Material), initial visualization of symbiont data from lab corals revealed differential symbiont community composition in the historical bleaching phenotypes, with historically non-bleached corals having mixed communities and historically bleached corals containing *Cladocopium* only. Changes in the concentration of *Cladocopium* symbionts over time appeared to vary between the two phenotypes. Thus, the concentration of *Cladocopium* symbionts over time was analyzed using a linear mixed effects model (package = *lme4*) with fixed effects of timepoint, historical bleaching phenotype, and temperature treatment, with random intercepts for fragments nested within colony. Additionally, the concentration of both *Cladocopium* and *Durusdinium* symbionts in the mixed community corals over time was analyzed using a linear mixed effects model with fixed effects of timepoint, symbiont genus, and temperature treatment, with random intercepts for fragments nested within colony. Data for both models were transformed using Tukey’s Ladder of Powers (package = *rcompanion*), and a post-hoc interaction analysis was conducted to determine statistical differences between factor levels (package = *emmeans*). Using the methods of Cunning, Silverstein, and Baker (2018), we related the photochemical efficiency of symbionts to their community composition. Regression coefficients for the proportion of *Durusdinium* symbionts against F_v_/F_m_ in mixed community corals were interpreted as the photochemical advantage of *Durusdinium* under ambient (at the first timepoint, before exposure to any thermal stress) and heated (at the final timepoint, after the peak water temperatures in the lab and the field had passed) conditions. For the analysis of heated corals, samples from corals in the lab and the field were used to increase power.

Normalized F_v_/F_m_ data (see the Supplementary Material) from the lab and field corals was analyzed using a linear mixed effects model with fixed effects of timepoint, colony, and temperature treatment, with random intercepts for colonies nested within historical phenotype. Historical phenotype was not included as a fixed effect due to violations of the homogeneity of variance assumption. A post-hoc interaction analysis was used to determine differences between factor levels. The photochemical efficiency of treatment corals during the natural heating event was of particular interest to us, so we further examined normalized F_v_/F_m_ in lab and field corals specifically at the thermal stress timepoint using a linear mixed effects model with fixed effects of treatment and historical bleaching phenotype, with random intercepts for colony and location of fragments in the lab or in the field. The significance of these random effects was explored using the *rand* function (package = *lmerTest*). A post-hoc interaction analysis was used to determine differences between factor levels.

In order to determine which characteristics of temperature profiles may be important for profile design, we explored the relative importance of total thermal stress accumulated versus temperature variation during treatments in coral response to the natural thermal stress event. We used linear regression to examine the relationship between the F_v_/F_m_ at the peak temperature stress timepoint and accumulated degree heating weeks (DHWs) during the temperature treatments. DHWs were approximated as the time spent above the 27.5°C maximum monthly mean (MMM) for Kāne’ohe Bay (NOAA/NDBC), with temperature stress beginning to accumulate at a bleaching threshold of MMM +1°C (28.5°C) (Wyatt et al. 2019). We compared the fit of this linear regression to a regression examining the relationship between the normalized F_v_/F_m_ and the mean hourly rate of change (ROC) in temperature in each temperature treatment (package = *TTR*). All analyses were performed in R Programming Language 3.6 (RStudio Team 2015) (package = *dplyr*).

## Results

### Symbiont community composition

Bleaching phenotypes observed during the 2015 bleaching event, where “bleached” refers to colonies that bleached and “non-bleached” refers to those that did not, continued to be distinguished by differential symbiont community composition. At the initial timepoint, all samples from historically non-bleached colonies contained both *Cladocopium* and *Durusdinium* symbionts, but were mostly dominated by symbionts of the genus *Durusdinium*, while none of the samples from historically bleached colonies contained detectable levels of the genus *Durusdinium* (Figure 2a). This differential symbiont community composition persisted throughout the temperature treatment, recovery, and natural heat stress periods (Figure 2b-d).

**Fig. 2.**
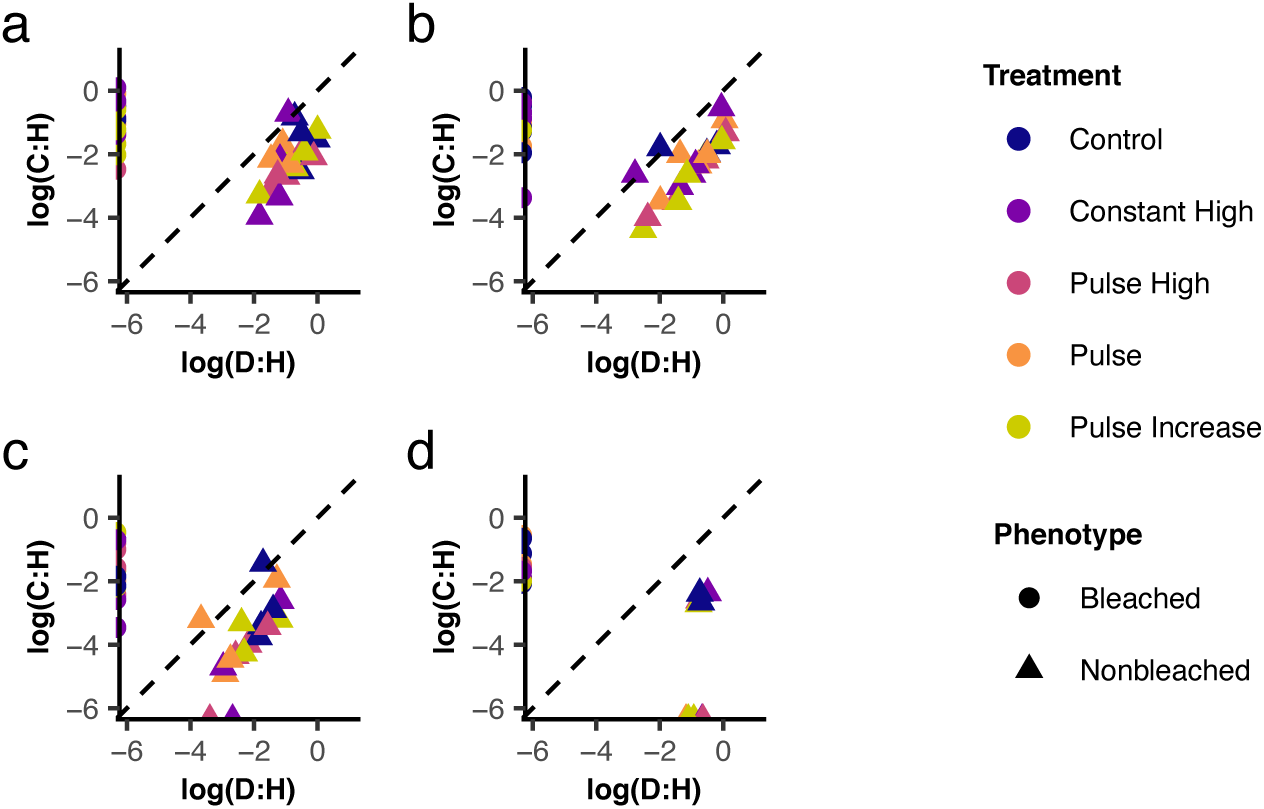
Base 10 log of *Cladocopium* to host cell (C:H) ratio against base 10 log of *Durusdinium* to host cell (D:H) ratio in samples in the lab at the four sampling timepoints: **(a)** pre-treatment (July 9^th^ 2019), **(b)** post-treatment (July 15^th^ 2019), **(c)** end of the recovery period (August 14^th^ 2019) and **(d)** peak of the natural thermal stress event (October 10^th^ 2019). Circles represent samples from colonies that bleached in 2015, triangles represent samples from colonies that did not. Colors indicate the five temperature treatments: control (blue), constant high (purple), pulse high (pink), pulse (orange), and pulse increase (yellow). Dashed diagonal lines represent a 50/50 ratio of *Cladocopium* to *Durusdinium* symbionts.

The lines formed by points of the mixed-community samples parallel to the diagonal line (x=y) in Figure 2 at the pre-treatment, post-treatment, and end of recovery timepoints indicate a stable ratio of *Cladocopium* to *Durusdinium* across mixed-community samples at these timepoints.

However, in the mixed-community colonies concentrations of *Cladocopium* decreased at the end of the recovery period (Figure 2c) and during the natural thermal stress event (Figure 2d), with some samples containing no detectable levels of *Cladocopium*.

### Changes in symbiont community composition during thermal stress

*Cladocopium*:host cell ratios were significantly higher in historically bleached corals than historically non-bleached, which also host *Durusdinium* (p<0.001; Figure 3a).There was a significant effect of time on the *Cladocopium*:host cell ratio across both phenotypes (p<0.001), where the mixed-community corals showed a significant decline in *Cladocopium*:host cell ratio during the recovery and natural thermal stress periods while corals hosting only *Cladocopium* did not (Figure 3a). In corals hosting both genera (i.e., non-bleached), there was a significantly higher concentration of *Durusdinium* than *Cladocopium* throughout the course of the experiment (p<0.001; Figure3b) and a significant effect of time (p<0.001), where symbiont concentrations significantly decreased following the temperature treatments (Figure 3b). However, there was also an interactive effect of symbiont and timepoint (p<0.001): during the natural thermal stress event *Cladocopium* concentrations continued to decline while *Durusdinium* concentrations recovered (Figure 3b).

**Fig. 3.**
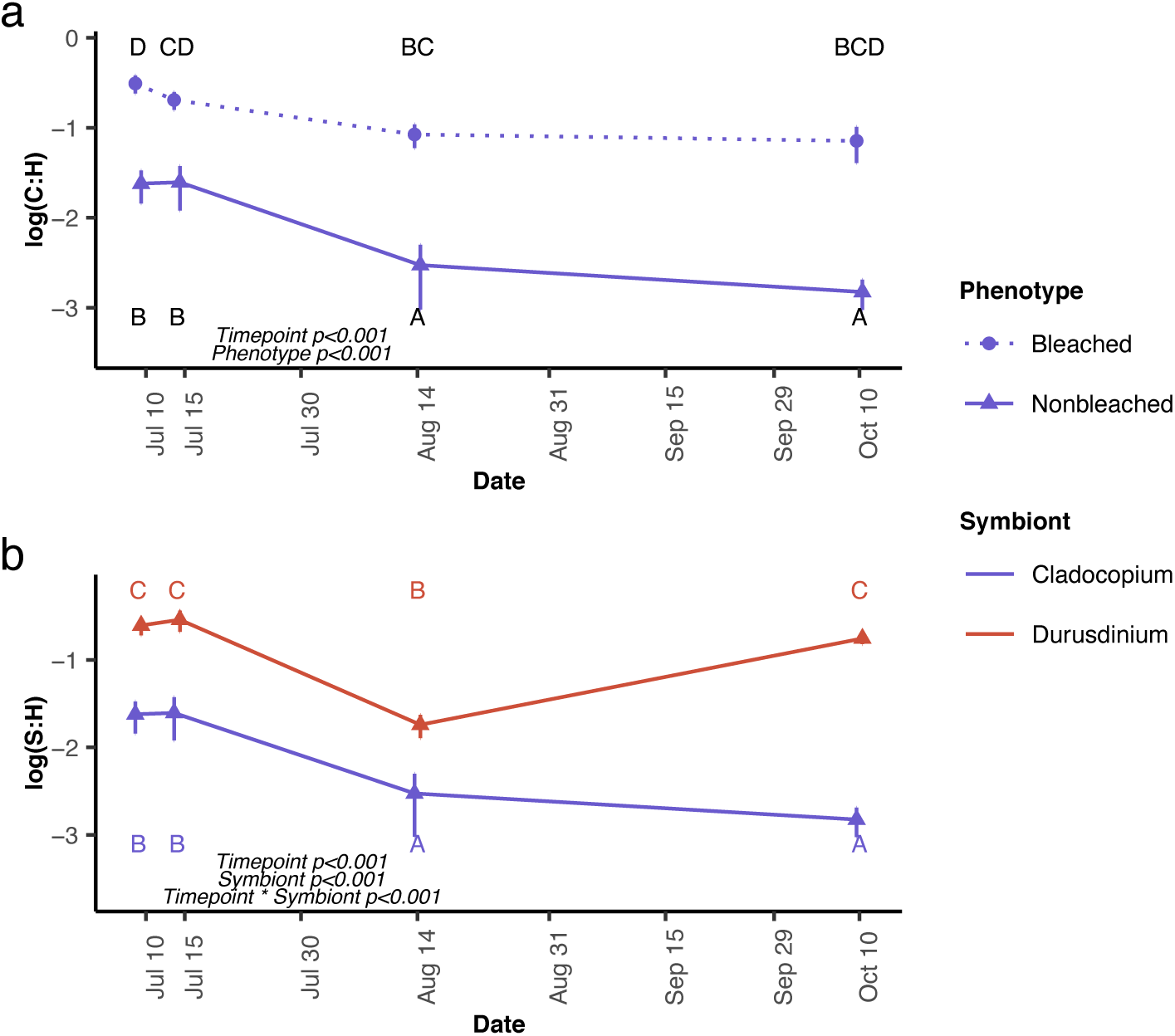
Mean base 10 log symbiont to host cell ratios of **(a)** *Cladocopium* symbionts in historically bleached, *Cladocopium*-only and non-bleached, mixed-community corals in the lab over time and **(b)** *Cladocopium* and *Durusdinium* symbionts in historically non-bleached corals in the lab over time. Error bars represent the standard error. Circles and dotted lines represent historically bleached corals, triangles and solid lines represent historically non-bleached corals. Blue indicates symbionts of the genus *Cladocopium*, red indicates symbionts of the genus *Durusdinium*.

### Photochemistry of symbiont communities

F_v_/F_m_ values were not significantly affected by the proportion of *Durusdinium* symbionts in mixed-community corals. Prior to thermal stress, there was no significant relationship between the proportion of *Durusdinium* and F_v_/F_m_ (p=0.41, slope=0.015, Figure 4a). At the final timepoint, when corals had experienced temperature treatments and a natural thermal stress event, there was a non-significant relationship between the proportion of *Durusdinium* and F_v_/F_m_ (p=0.36, slope=-0.0085, Figure 4b).

**Fig. 4.**
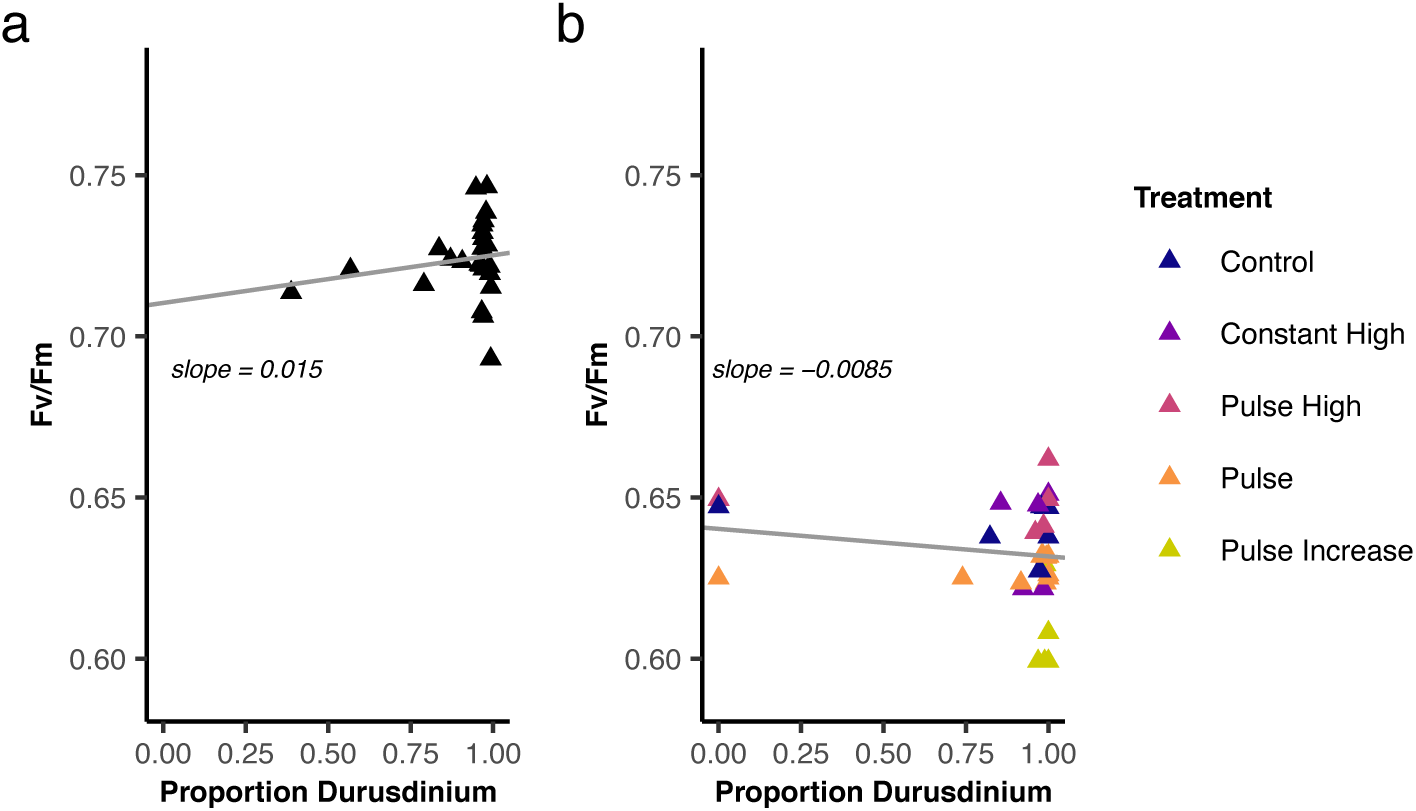
Linear regression plots between mean colony F_v_/F_m_ and proportion *Durusdinium* in mixed-community samples at **(a)** the initial timepoint, before exposure to thermal stress, and **(b)** the final timepoint, after temperature treatments and the natural thermal stress event. Colors indicate the five temperature treatments: control (blue), constant high (purple), pulse high (pink), pulse (orange), and pulse increase (yellow).

### Effects of genotype and treatment on photochemical efficiency

There was a significant main effect of temperature treatment (p<0.001) and timepoint (p<0.001), as well as an interactive effect of treatment and timepoint (p<0.001) on photochemical efficiency. After the temperature treatments, corals in all treatments had significantly lower normalized F_v_/F_m_ compared to the control (Figure 5a). After the recovery period, normalized F_v_/F_m_ across all treatments converged, but following the natural thermal stress event corals in the control and constant high treatments had significantly higher normalized F_v_/F_m_ than those in the various pulsed treatments (Figure 5a). There was also a significant interactive effect of treatment and genotype on normalized F_v_/F_m_ (p=0.048), with treatment effects on photochemical efficiency varying significantly by genotype (Figure S1) even between paired colonies with shared environmental histories (Figure 5b).

**Fig. 5.**
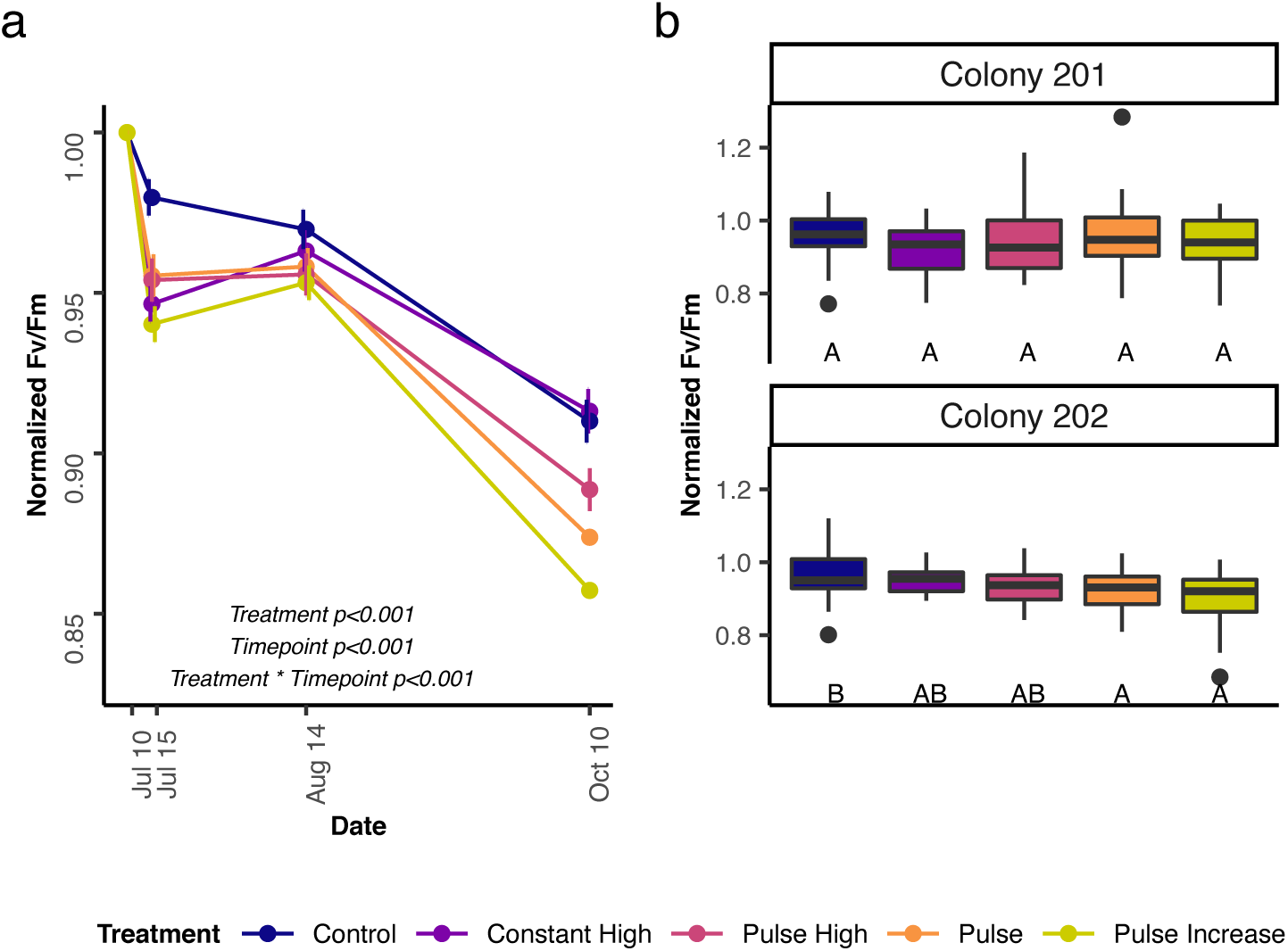
**(a)** Mean normalized F_v_/F_m_ by temperature treatment over time. Error bars represent the standard error. **(b)** Boxplots of normalized F_v_/F_m_ by treatment averaged over time in paired colonies 201 (historically bleached) and 202 (historically non-bleached), with the middle line showing mean normalized F_v_/F_m_. Letters represent the differences between treatment groups found during post-hoc interaction analysis. Colors indicate the five treatments: control (blue), constant high (purple), pulse high (pink), pulse (orange), and pulse increase (yellow).

### Photochemical efficiency in the natural thermal stress event

At the peak of the natural thermal stress event, there was a significant effect of historical bleaching phenotype (p=0.012) and treatment (p<0.001), but no significant effect of colony (p=0.33) on photochemical efficiency. Historically bleached corals had significantly higher normalized F_v_/F_m_ than historically non-bleached corals (Figure 6a). None of the treatments were associated with significantly higher photochemical efficiency than the control (Figure 6a).

**Fig. 6.**
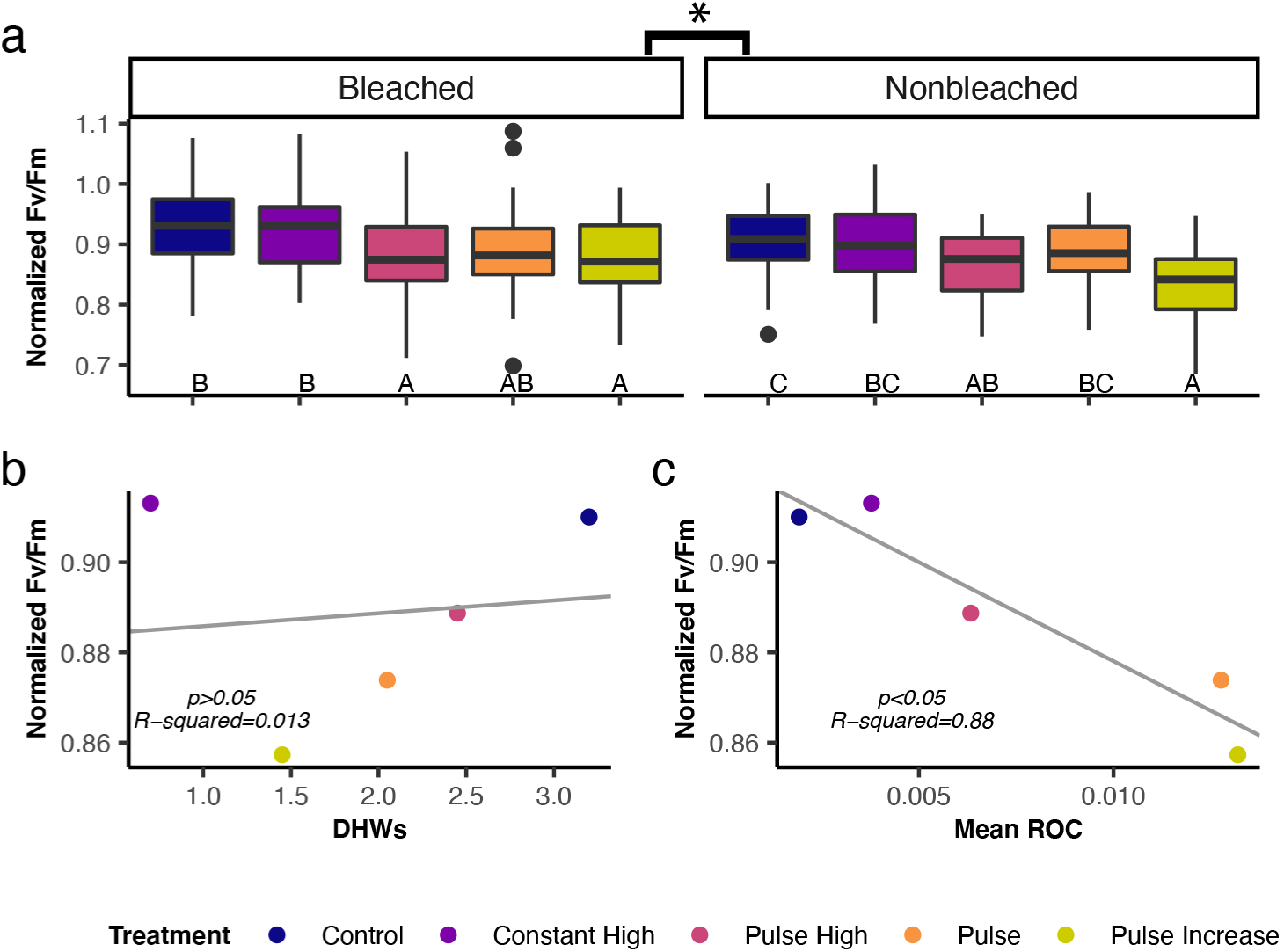
**(a)** Boxplot of normalized F_v_/F_m_ at the peak thermal stress timepoint in the historical bleaching phenotypes by treatment, with the middle line showing the mean normalized F_v_/F_m_. Letters represent the differences between treatment groups found during post-hoc interaction analysis. Linear regression plots between normalized mean F_v_/F_m_ by treatment at the peak of the thermal stress event and **(b)** the accumulated degree heating weeks (DHWs) during acclimatization treatments and **(c)** the mean hourly rate of change in temperature during the acclimatization treatments. Colors indicate the five acclimatization treatments: control (blue), constant high (purple), pulse high (pink), pulse (orange), and pulse increase (yellow).

However, in the historically bleached phenotype, corals that were exposed to the constant high treatment had significantly higher normalized F_v_/F_m_ than those in the pulse high or pulse increase treatments, though they were not significantly different from the pulse treatment (Figure 6a). In the historically non-bleached phenotype, photochemical efficiency in constant high corals was not significantly different from the pulse or pulse high treatments (Figure 6a). There was no significant relationship between the number of degree heating weeks (DHWs) accumulated during temperature treatments and the mean normalized F_v_/F_m_ of corals in these treatments at the peak of the natural thermal stress event (p=0.85, Figure 6b). However, there was a strong significant negative relationship between the mean hourly rate of change (ROC) during the temperature treatments and the mean normalized F_v_/F_m_ of corals in the corresponding treatments (p=0.012, Figure 6c).

## Discussion

The results demonstrate that historical bleaching phenotypes in *M. capitata* in Kāne’ohe Bay are underpinned by stable differences in symbiont community composition, which continued to differentiate the phenotypes throughout exposure to thermal stress in this study. *Cladocopium*-only corals that bleached in 2015 lost less *Cladocopium* symbionts than the mixed-community corals that did not, which preferentially recovered *Durusdinium* during the natural warming event. Additionally, response to the temperature treatments was modulated by host genotype, and although the profiles did not elicit protective acclimatization, temperature variation in our treatments had a significant detrimental effect on coral response to the natural thermal stress event.

### Variation in symbiont community composition

The connection between bleaching phenotype during the 2015 bleaching event and dominant symbiont genus observed by Cunning and colleagues (2016) persisted in this study: samples taken from historically non-bleached colonies still contained mixed communities of *Cladocopium* and *Durusdinium* that were mostly dominated by thermally tolerant *Durusdinium*, while historically bleached colonies contained only *Cladocopium*, although it is possible that these samples contained low concentrations of *Durusdinium* that were not detected through qPCR (Mieog et al. 2009). While previous work has established that some coral species with multi-partner symbioses can expand their thermal tolerance by “shuffling” their symbionts to change the dominant symbiont genus (Baker 2003; Silverstein et al. 2014; Cunning et al. 2015), neither the multi-year bleaching events in 2014 and 2015 (Bahr et al. 2017) nor the temperature treatments and natural thermal stress event in this study appeared to induce symbiont shuffling in *M. capitata*. Mixed-community colonies maintained *Durusdinium*-dominated symbiont communities four years post-bleaching despite the potential for a competitive advantage of *Cladocopium* under ambient conditions (McIlroy et al. 2019). Similarly, different *M. capitata* genotypes with mixed communities of algal symbionts had remarkably similar *Cladocopium* to *Durusdinium* ratios. This suggests that the differences in symbiont community composition underlying differential bleaching phenotypes during the 2015 event are highly stable, similarly to symbiont communities in *Pocillopora* spp. in the eastern Pacific (McGinley et al. 2012). These stable differences in colonies immediately adjacent to each other suggest the coral host plays an important role in modulating differences in symbiont community composition, possibly through the host immune system (Roach et al. 2020). However, additional work has also shown that not all *Cladocopium*-dominated *M. capitata* colonies bleached in 2015, indicating that symbiont identity is not the sole factor driving historical bleaching phenotype (Cunning et al. 2016; Ritson-Williams and Gates 2020).

The differences in symbiont community composition between the historical bleaching phenotypes were also evident in the differential loss of *Cladocopium* symbionts: following the temperature treatments and during the natural thermal stress event, mixed-community corals exhibited greater losses of *Cladocopium* symbionts compared to *Cladocopium*-only corals.

Additionally, in the months following the temperature treatments, mixed-community corals recovered *Durusdinium*, but continued to lose *Cladocopium*, indicating some plasticity in the symbiont community of mixed-community corals. However, these changes are not particularly strong and may not strongly impact thermal tolerance: Cunning and colleagues (2016) found that symbiont shuffling in *M. capitata* in favor of *Durusdinium* following thermal stress was very low, and also found an instance of community dominance switching to the more thermally sensitive *Cladocopium*.

The trends in the loss of *Cladocopium* from mixed-community corals may indicate some competitive advantage of *Durusdinium*. Previous studies have found that *Durusdinium* is able to outcompete *Cladocopium* only in thermally stressful conditions (McIlroy et al. 2019), or when tradeoffs regarding growth (Jones and Berkelmans 2010; Cunning et al. 2015), nutrient acquisition (Baker et al. 2013), and photochemical efficiency (Cantin et al. 2009) diminish. Thus, the conditions of thermal stress during the mild natural warming event may have been enough to promote the preferred persistence of *Durusdinium* in mixed symbiont communities, but still mild enough for *Cladocopium* to perform relatively well in corals with no competition with *Durusdinium*. The significantly lower *Cladocopium*:host cell ratio in mixed-community corals at the end of the thermal stress event may also explain why these historically non-bleached corals were less photochemically efficient at this timepoint, as symbionts of the genus *Cladocopium* have been shown to have greater relative electron transfer rates through photosystem II at ambient temperatures than their *Durusdinium* counterparts (Cantin et al. 2009). Additionally, the tendency of mixed-community corals to preferentially recover *Durusdinium* over *Cladocopium* may also contribute to the stable differences in symbiont community composition between the historical bleaching phenotypes observed in this study.

This stability in symbiont community composition may also arise from a lack of photochemical advantage of one symbiont type over the other. In a 2018 study, Cunning and colleagues found that differences in symbiont shuffling between species were strongly related to differences in photochemical efficiency of co-occurring symbionts during times of thermal stress. In the mixed-community *M. capitata* colonies in this study, no significant photochemical advantage of *Cladocopium* or *Durusdinium* symbionts was observed under ambient or thermally stressful conditions, possibly contributing to the lack of symbiont shuffling as a response to thermal stress.

### Influence of host genotype

Although the intention of the temperature treatments was to expose coral fragments to protective levels of thermal stress, symbiont photochemical efficiency decreased significantly in all treatments compared to the control, indicating damage to the photosynthetic apparatus of the algal symbionts (Warner 1999; Jones et al. 2000). The response to the temperature treatments over time varied significantly by genotype, even amongst neighboring colonies with the same environmental histories, indicating a significant role of coral host genotype in the response to thermal stress. Notably, this genotypic variation included higher photochemical efficiency in constant high fragments compared to other temperature treatments in colony 12 during natural thermal stress (Figure S1), suggesting host genotype could play a role in modulating the efficacy of these treatments. Past work has shown that there is a strong genetic basis of thermal tolerance in the coral host (Dixon et al. 2015), and that genotypic and environmental differences underlie the differential bleaching responses of corals hosting the same symbiont type (Drury et al. 2017). As the *M. capitata* colonies in this study are characterized by stable symbiont community differences, variable response of different host genotypes to temperature treatments suggests that changes in host gene expression could play a more important role in the acclimatization of *M. capitata* than mechanisms such as symbiont shuffling (Bellantuono et al. 2012b; Bay and Palumbi 2015; Ainsworth et al. 2016).

### The role of temperature treatments

Despite the photochemical damage caused during the temperature treatments, a mild thermal stress event in Kāne’ohe Bay allowed us to explore the effects of the treatments and their interactions with the differences in symbiont community underlying previous bleaching phenotype in shaping the response of *M. capitata* to natural thermal stress. Although none of the treatments had a protective effect during the thermal stress event, fragments that had been in the constant high treatment were more photochemically efficient at the peak of thermal stress than those that experienced one of the three pulsed treatments, though they did not perform better than controls. Differences in photochemical efficiency between the treatments appeared to arise from differences in temperature variability, rather than the amount of thermal stress accumulated: pulsed treatments, which had more variation in temperature, had a more detrimental effect than the constant temperatures that characterized the constant high and control treatments. This result contradicts a previous study by Bay and Palumbi (2015) which found that exposure to pulsed temperature profiles conferred thermal tolerance while a constant temperature treatment did not. However, past work has also documented that diel variations in temperature can elicit a detrimental response in reef-building corals (Putnam and Edmunds 2010). Additionally, it is possible that our constant high treatment did not expose corals to enough thermal stress to cause them to deteriorate significantly, obscuring the benefits of the intermittent recovery during pulsed treatments. In the constant high treatment, corals were exposed to approximately 3.1 DHWs, which is well below the 4 DHWs usually assumed to induce coral bleaching (NOAA/NESDIS). However, work by Ruiz-Jones and Palumbi (2017) showed that fine-scale adjustments in gene expression occur at temperatures well below the bleaching threshold, suggesting these mechanisms may have been overwhelmed by the high, significantly fluctuating temperatures in our treatments. Thus, further investigation into the design of temperature profiles intended to induce acclimatization should consider both the overall accumulation of thermal stress as well as the variation in temperature by comparing the effects of acute, high temperature profiles such as the ones in this study to those of extended treatments at lower temperatures.

The experimental treatments used here appeared to be more damaging than protective overall, though this response varied by genotype, which makes it difficult to draw conclusions regarding the utility of these profiles in inducing an acclimatization response. Hawaiian reef corals experience stress and decrease skeletal growth at temperatures above 27°C (Bahr et al. 2015b). Although temperature tolerance for *M. capitata* in Kāne’ohe Bay can be higher than this threshold (Jury and Toonen 2019), ambient temperatures in our control tanks during the treatments periodically exceeded the Kāne’ohe Bay bleaching threshold of 28.5°C (NOAA/NDBC, Wyatt et al. 2019). Thus, it is likely that corals were already experiencing thermal stress under “ambient” conditions, and that the addition of further stress through the temperature profiles caused photochemical damage, obscuring any possible benefit that may have been derived from the profiles. This is supported by the fact that corals in the control tanks also saw decreases in photochemical efficiency during the treatment and recovery periods. This underlines the importance of understanding how long acclimatization effects can persist after treatment, as our results indicate that exposure to increased temperatures may need to occur well before earlier seasonal warming under climate change in order to be effective in inducing a protective acclimatization response. This finding is supported by the work of Ainsworth et al. (2016), which found that increased temperatures due to climate change had a disabling effect on reconstructed natural protective temperature trajectories recreated in a laboratory setting.

### Implications for acclimatization

These results indicate that artificial protective temperature treatments should consider the interactions between acute, high temperature treatments and extended treatments at lower temperatures. Additionally, given the potential cumulative stressors of increasing sea surface temperatures and lab-based temperature treatments, the timing of these treatments and their persistence over longer periods of time needs to be better understood. Finally, we suggest that in corals with stable symbiont communities that do not “shuffle” their symbionts to increase thermal tolerance, such as the *M. capitata* colonies in this study, varying thermal tolerance arising from genotypic differences in the coral host and changes in the host’s gene expression will play a significant role in successful acclimatization to thermal stress.

## Author Contributions

**JD, CD**, and **CC** conceptualized the experiment. **JD, VAK, CD**, and **CC** collected data. **JD** analyzed data with assistance from **CD** and **ACB. JD, CD** and **ACB** interpreted the data. **JD** wrote the manuscript. All authors edited and approved the final version.

## Acknowledgements

We thank Nina Bean, Luke Kikukawa, Shayle Matsuda, Josh Hancock, Joel Huckeba, Ariana Huffmyer, Ty Roach, Mariana Rocha de Souza, and the Gates Coral Lab for their assistance with this project. This work was funded by the Paul G. Allen Family Foundation and formed part of JD’s Master’s thesis at the University of Miami.

## Competing Interests

We declare no competing interests.

## Data availability

All raw data and code are available on Zenodo at https://doi.org/10.5281/zenodo.3946512

## Notes

https://doi.org/10.5281/zenodo.3946512

### Competing Interest Statement

The authors have declared no competing interest.

