## Supplementary Figure 1 and Supplementary Materials and Methods for "Host genotype and stable differences in algal symbiont communities explain patterns of thermal stress response of *Montipora capitata* following thermal pre-exposure and across multiple bleaching events"

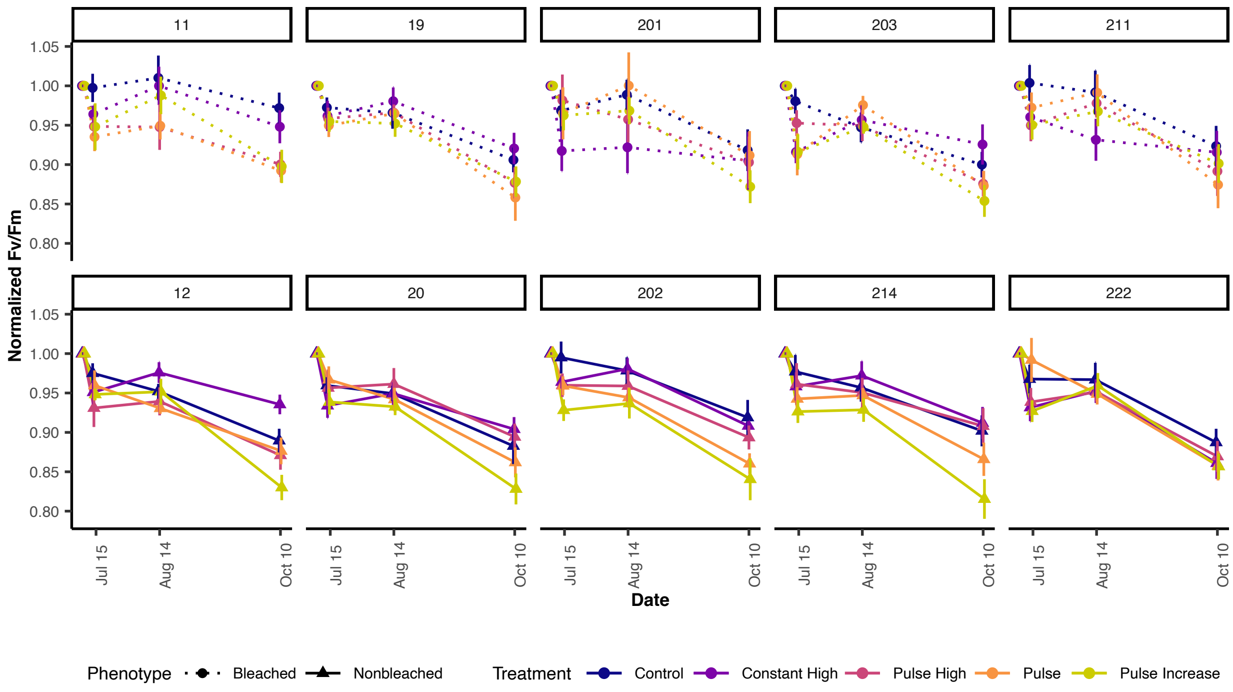


**Fig. S1.** Mean normalized $F_v/F_m$ by temperature treatment in each colony over time. Circles and dotted lines represent colonies that bleached in 2015, triangles and solid lines represent colonies that did not bleach in 2015. Colors indicate the five temperature treatments: control (blue), constant high (purple), pulse high (pink), pulse (orange), and pulse increase (yellow). Error bars represent the standard error.

**Supplementary Methods and Materials:**

*Design of temperature treatments*

In order to investigate whether short, acute artificial temperature treatments that would be practical to administer in management and restoration settings could be used to induce acclimatization to thermal stress, we designed four different temperature treatments with high maximum temperatures. In order to determine the appropriate maximum temperature of the profiles, *M. capitata* fragments of opportunity were ramped up to varying peak temperatures over the course of 16 hours, then kept at this temperature for six days. Fragments were exposed to maximum temperatures of 30.5°C, 31°C, 31.5°C, 32°C, and 32.5°C, and photographed daily to track bleaching response. After the six-day trial, corals in the 32°C and higher temperature treatments had visibly bleached. Thus, 31.5°C was chosen as the maximum temperature for the acclimatization profiles in order to ensure coral fragments in all treatments, including constant high temperature conditions, would be exposed to sublethal thermal stress.

*Data cleanup and normalization*

Following visualization of the distribution of PAM fluorometry data, $F_v/F_m$ values less than 0.52 were removed from the dataset, as lower densities of light-absorbing Symbiodiniaceae in the tissues of thermally stressed corals can cause issues with PAM fluorometry measurements due to backscattering of light off the coral skeleton (Wangpraseurt et al. 2019). $F_v/F_m$ values higher than

0.84 were also removed from the dataset, as this is the theoretical maximum $F_v/F_m$ measurement using PAM fluorometry (Walz, 2020). To account for colony and fragment differences at the start of the experiment, $F_v/F_m$ values at the post-treatment, recovery, and peak thermal stress timepoints were normalized by fragment by dividing by the initial pre-treatment $F_v/F_m$. qPCR runs with coral host $C_T$ values greater than the $75^{th}$ percentile plus 1.5 times the interquartile range were removed from the dataset, as high host $C_T$ values may indicate low DNA yield or carryover of PCR inhibitors during extraction (Cunning et al. 2016 Supplementary Materials). qPCR data from one fragment from colony 11 was excluded from analysis, as samples from this fragment demonstrated inconsistencies in symbiont types detected between timepoints.

*Reference List*


Cunning R, Ritson-Williams R, Gates RD (2016) Patterns of bleaching and recovery of Montipora capitata in Kaneohe Bay, Hawaii, USA. Mar Ecol Prog Ser 551: 131-139

Wangpraseurt D, Lichtenberg M, Jacques SL, Larkum AWD, Kühl M (2019) Optical properties of corals distort variable chlorophyll fluorescence measurements. Plant Physiol 179: 1608-1619

Walz (2020) Diving-PAM-II: Manual for standalone use. Effeltrich, Germany
